# Transcriptomic analysis identifies injury-responsive fibroblast populations as potential mediators of Wnt-dependent spinal cord regeneration

**DOI:** 10.1101/2024.05.17.594712

**Authors:** Samuel R. Alper, Deeptha Vasudevan, Maya K. Wheeler, Samin Panahi, Richard I. Dorsky

## Abstract

**Background:** In humans and other mammals, spinal cord injury (SCI) can lead to a permanent loss of sensory and motor function, due to the inability of damaged neurons and axons to regenerate. However, other vertebrate species including zebrafish exhibit complete spinal cord regeneration and functional recovery after SCI. Wnt signaling is required for neurogenesis and axon regrowth in a larval zebrafish SCI model, but the genes regulated by this pathway and the cell types that express them remain largely unknown.

**Results:** In this study, we used bulk RNA-sequencing (RNAseq) to identify candidate genes regulated by Wnt signaling that are expressed after SCI. Using this unbiased screen, we identified multiple genes previously unassociated with SCI in larval zebrafish, and confirmed by in situ hybridization that their expression is injury-responsive, Wnt-dependent, and localized to fibroblast-like cells surrounding the spinal cord.

**Conclusions:** Together, our data reveal potential novel gene targets and cell populations that may play important roles in spinal cord regeneration.

## INTRODUCTION

Treatment options for human spinal cord injury (SCI) are limited and no current therapies promote full functional recovery. Permanent sensorimotor deficits following SCI in the adult mammalian spinal cord are due to the absence of progenitor cells that can replace lost neurons ^1^, intrinsic deficits in the capacity for axon regrowth by surviving neurons ^2^, as well as the demonstrated extrinsic actions of surrounding cells. Several cell types including glia, immune cells, mesenchymal fibroblasts, and meningeal cells infiltrate the injury site, induce inflammation, and secrete an array of growth-inhibiting extracellular matrix (ECM) components ^3^. The result is the formation of a fibrotic scar that poses both a molecular and physical barrier for repair of the injury site. However, the degree of spinal cord regeneration varies widely between different vertebrate classes, suggesting that the mechanisms underlying functional recovery in spontaneously regenerating species could lead to future therapeutic approaches in humans. Zebrafish have been developed as a model system for studying SCI because they exhibit regenerative neurogenesis, axon regrowth, and rewiring of spinal circuits following SCI, with eventual functional recovery ^4-7^. The larval zebrafish SCI model is particularly advantageous due to its high throughput, rapid regeneration time, optical accessibility, and general conservation of regenerative mechanisms with adult fish ^8^. Using this model, our goal is to identify the mechanisms that promote spinal cord regeneration, and determine whether they are evolutionarily conserved.

A wide array of cell types and signaling cascades are activated following zebrafish SCI. Of particular importance is the injury-induced reactivation of signaling pathways that play prominent roles during embryonic development, including the Wnt/β-catenin pathway ^9^. After SCI in larvae, Wnt signaling activity begins within 1 day post-injury (dpi), peaks by 2 dpi, and dissipates by around 3 dpi ^10^. Prior data from our lab and others shows that Wnt signaling is required for both regenerative neurogenesis and axon regrowth ^6,10-12^. After SCI at both larval and adult stages, resident neural progenitors proliferate and differentiate into newborn neurons. However, only a small fraction of spinal cord progenitors and no surviving neurons appear to respond directly to Wnt signaling after SCI ^6,10^.

Instead, most Wnt-responsive cells after injury are extraspinal and express general markers of fibroblasts, but more specific identification and characterization of these populations has remained elusive ^10^. Wnt pathway activity in fibroblast-like cells after SCI is required for the expression of collagens and other ECM components. However, so far only two genes encoding collagen isoforms have been shown to be induced by SCI, Wnt-dependent, and required for axon regrowth ^10^. While valuable, these studies have not yet led to a complete understanding of the Wnt target genes that mediate spinal cord regeneration and the identity of extraspinal Wnt-responsive cell populations that express them.

To address these questions, we performed bulk RNA sequencing (RNAseq) analyses on injured zebrafish larvae treated with pharmacological inhibitor of Wnt signaling. By comparing mRNA expression from these larvae with untreated controls, we have identified Wnt-dependent genes that we have been able to functionally categorize based on gene ontology (GO) enrichment analysis. Of particular interest are genes encoding feedback inhibitors of Wnt signaling, and markers of injury-associated infiltrating and meningeal fibroblasts. We have validated the SCI-and Wnt-dependent expression patterns of transcripts from each of these categories in the regenerating larval spinal cord by in situ hybridization. Together, our data identify novel Wnt target genes and cell populations that may play important roles in spinal cord regeneration.

## EXPERIMENTAL PROCEDURES

### Zebrafish

The following zebrafish strains were used for this study: *AB, *mitfa*^*w2* 13^, and *Tg(7xTCF-Xla*.*Siam:GFP)*^*ia4* 14^. Larvae used for experiments were obtained from incrosses and staged according to ^15^. All experiments were approved by the University of Utah Institutional Animal Care and Use Committee.

### Bulk RNA sequencing

*AB and *Tg(7xTCF-Xla*.*Siam:GFP)*^*ia4*^ larvae were raised in E3 media until 5 dpf (days post fertilization), anesthetized in 0.016% Tricaine (Sigma), and injured by complete transection of the spinal cord at the level of the anal pore using a sharpened and sterilized FST injury tool. Larvae were allowed to recover for 6 hours in E3 before being transferred into media containing E3 with 35μM IWR-1 (Sigma) in 0.25% DMSO or 0.25% DMSO as a vehicle control. Larvae were fed rotifers starting at 1 dpi until fixation at 3 dpi (8 dpf). To confirm Wnt pathway inhibition, *Tg(7xTCF-Xla*.*Siam:GFP)*^*ia4*^ larvae were imaged with a Nikon A1 HD25 confocal microscope using a 60X oil immersion lens and images were processed in Adobe Photoshop.

Larvae were fixed in 4% paraformaldehyde (PFA) and 10% sucrose in PBS at 3 dpi at 4°C overnight. For each biological replicate of 20 animals, spinal cord and adjacent tissues dorsal to the notochord were dissected 1-2 segments caudal and rostral to the injury site. This process was repeated for three biological replicates per treatment condition. The Invitrogen Recover All Total Nucleic Acid Isolation Kit for FFPE was used to extract RNA from the tissue samples.

Purified RNA samples were analyzed by the High Throughput Genomics Core Facility at the Huntsman Cancer Institute using the NEBNext Ultra II Directional RNA Library Prep Kit with rRNA depletion. cDNA libraries were then sequenced using a NovaSeq 6000 instrument at 50X50 bp read length and 100M read-pair increments. Cutadapt was used for adapter trimming, FastQC for quality control, STAR for alignment and featureCounts for quantification with genome and annotation downloaded from Ensembl version Danio_rerio.GRCz11.94, and DESeq2 for differential expression analysis. Gene Ontology (GO) enrichment analysis for biological process and cellular component was performed for all genes with a negative log2 fold change and adjusted p values <0.05. Sequencing data was submitted to GEO under accession #GSE274820.

### In situ hybridization and immunohistochemistry

*mtfa*^*w2*^ homozygous mutant larvae were raised in E3 media until 3 dpf, anesthetized in 0.016% Tricaine (Sigma), and injured by complete transection of the spinal cord at the level of the anal pore using a sterile 30G needle ^16^. Injured larvae were allowed to recover in E3 for 6 hours before being transferred to media containing either 15uM IWR-1 in 0.25% DMSO or 0.25% DMSO as a vehicle control ^10^. In situ hybridization probes were made by a clone-free method as described previously^17^with DNA templates purified using Zymo Research DNA clean & concentrator^TM^-5 kit. Primers were designed by Primer-BLAST (NCBI).

Injured fish were fixed at 2 dpi in 4% PFA containing 0.1% Tween 20 for 1 hour at room temperature and subsequently dehydrated in 100% Methanol and stored at -20°C. Whole mount *in situ* hybridization and IHC were performed according to ^16^ with a 10-minute pre-incubation in 2%H_2_O_2_ in MeOH. ISH-stained fish were imaged on an Olympus compound microscope using a 20X objective. Chicken anti-GFP (Aves #GFP-2010) and Goat anti-Chicken Alexa-488 (Invitrogen #A-11039) antibodies were used to label GFP+ cells in *Tg(7xTCF-Xla*.*Siam:GFP)*^*ia4*^ larvae. 30um Z-stacks of antibody-labeled larvae were obtained using a Zeiss700 confocal microscope with a 20x objective. Linear adjustments to brightness and contrast were made using FIJI.

## RESULTS AND DISCUSSION

### RNAseq analysis reveals known and novel Wnt-dependent genes following SCI

Previous work has demonstrated that Wnt signaling is integral to both axon regrowth and neurogenesis after injury ^6,10^. However, the transcriptional targets of Wnt signaling that mediate these regenerative processes remain incompletely characterized. To generate a list of candidate target genes, we performed bulk RNAseq on regenerating spinal cord and adjacent tissues dissected from larvae treated with the Wnt pathway inhibitor IWR-1 ^18^, and from vehicle-treated controls. Tissue was collected at 3 dpi following spinal cord transections performed on 5 dpf larvae (Fig. 1A). Previous work from our lab found that at this recovery timepoint axon regrowth is observed at the injury site, but regenerated neurons are not yet present ^6,7^. After DESeq2 analysis of IWR-1 treated vs. control samples, we were able to identify 1,791 differentially expressed annotated genes.

**Figure 1.**
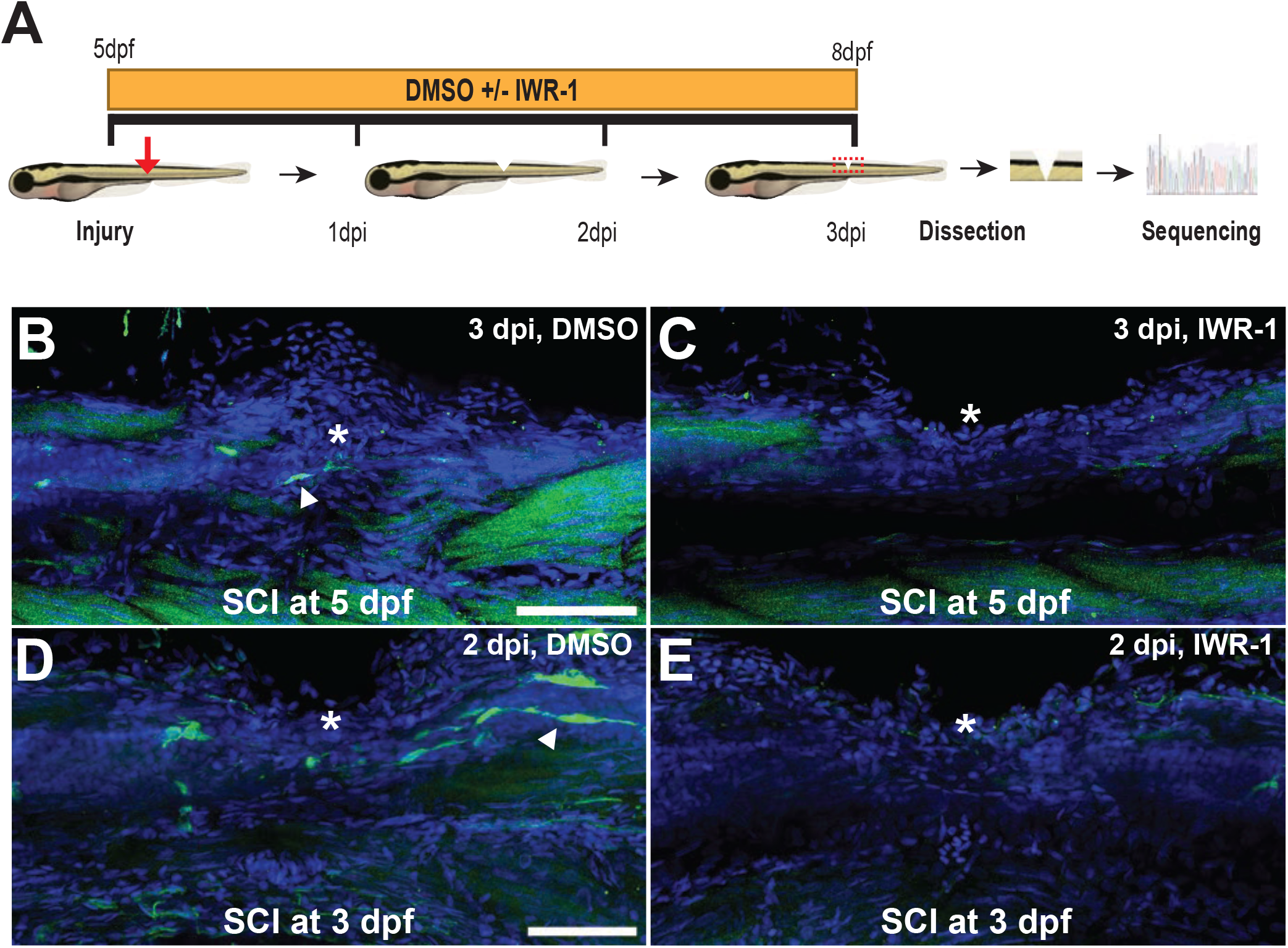
Workflow for bulk RNA sequencing and validation of Wnt inhibition by IWR-1. (A) Schematic of RNAseq workflow. 5 dpf larvae were injured and treated with IWR-1 or vehicle control (DMSO) as a vehicle control. At 3 dpi, a 1mm segment of tissue including the injury site and surrounding tissue was extracted, dissociated, and sent for bulk RNAseq. (B-C) SCI was performed on *7xTCF-Xla*.*Siam:GFP* fish at 5 dpf and treated with IWR-1 or DMSO until fixation at 3 dpi. Wnt-responsive cells (GFP+) are present at the injury site both within and dorsal to the spinal cord in vehicle-treated controls. A decrease in Wnt-responsive cells in IWR-1 treated animals indicates successful pharmacological inhibition of Wnt signaling. (D-E) SCI was performed on 3 dpf WT larvae and treated continuously with either IWR-1 or DMSO until sacrifice at 2 dpi. Wnt-responsive (GFP+) cells are present at the injury site in DMSO controls. GFP expression is reduced in IWR-1 treated fish. At both timepoints, analysis of GFP expression in the transgenic Wnt reporter line shows successful IWR-1 mediated inhibition of Wnt signaling. Asterisk: Injury site.

Because we are primarily interested in genes with expression that is induced by SCI, and because most known direct Wnt/ß-catenin targets are transcriptionally activated by pathway activity ^19^, we focused specifically on 872 genes with expression that was significantly decreased by Wnt inhibition (adjP<0.05). When we performed a gene ontology (GO) analysis of this group using the PANTHER overrepresentation test for biological process, we found significant enrichment of “negative regulators of Wnt signaling pathway” (13.5-fold enrichment, adjP=3.24E-04), which included multiple genes that are known direct feedback targets of Wnt signaling. This result confirmed the effectiveness of our screen in detecting Wnt-dependent gene expression. We also found significant enrichment of GO terms related to tissue development and regeneration, suggesting that some novel Wnt targets identified by our screen may be involved in the injury response. When we performed a GO analysis using the PANTHER overrepresentation test for cellular component, we found high enrichment of “collagen-containing extracellular matrix” (5.02-fold enrichment, adjP=4.57E-03), which was consistent with the known requirement of Wnt-dependent Collagen XII expression by infiltrating fibroblasts for axon regrowth after SCI ^10^ (Fig. 3A). Together, these data suggested that our RNAseq analysis could help identify novel Wnt-dependent molecular and cellular mediators of spinal cord regeneration.

### Validation of candidate Wnt target gene expression in a 3 dpf injury model

While our RNAseq screen was performed after SCI at 5 dpf, several recent studies have adopted an earlier 3 dpf injury paradigm for modeling spinal cord regeneration ^10,20,21^. Following injury at the 3 dpf timepoint, the zebrafish larva is more accessible for subsequent whole-mount imaging and histological analysis, and the regenerative response largely recapitulates the responses in both 5 dpf larval and adult injury paradigms ^8^. In order to confirm that Wnt signaling was similarly injury-responsive and IWR-1-dependent following SCI at both 3 and 5 dpf, we examined *Tg(7xTCF-Xla*.*Siam:GFP)*^*ia4*^ larvae, which express a GFP reporter of Wnt/ß-catenin pathway activity ^14^. Using both injury paradigms, we observed multiple GFP+ cells near the injury site in control larvae, while IWR-1 treated larvae showed a clear reduction in GFP expression (Fig. 1B-E). Based on these results we decided to use the 3 dpf injury model to validate the injury-responsive and Wnt-dependent expression of candidate genes from our RNAseq analysis.

### Injury-dependent expression of Wnt signaling feedback inhibitors

Our RNAseq analysis identified several known Wnt target genes encoding secreted inhibitors of pathway activity, including orthologs of *notum1, sfrp1, wif1, and dkk1* (Fig. 2A). We performed in situ hybridization for *notum1a* mRNA to determine whether this gene is expressed at the injury site following SCI. At 2 dpi, *notum1a* expression was present in close proximity to the injury site in vehicle-treated controls, but was clearly diminished in IWR-1 treated larvae (Fig. 2B-C’). This experiment confirmed the injury and Wnt dependence of a candidate identified by our screen, the efficacy of our drug treatment, and suggested that *notum1a* may play a currently unexplored role in Wnt-dependent spinal cord regeneration. While the requirement for the injury-induced activation of Wnt signaling in zebrafish spinal cord regeneration is well documented ^6,10,12^, other roles of Wnt signaling also require the subsequent down-regulation of pathway activity in order for complete function to be achieved ^22,23^. Studies have also shown that transient cell-type specific signals, including the Wnt pathway, allow cellular populations to execute pro-regenerative functions ^20,21,24-26^. Consistent with this possibility, secreted inhibitors such as Notum1 may act to restrict Wnt-signaling from cells near the injury site. Future work will be required to determine the precise dynamics of Wnt signaling after SCI.

**Figure 2.**
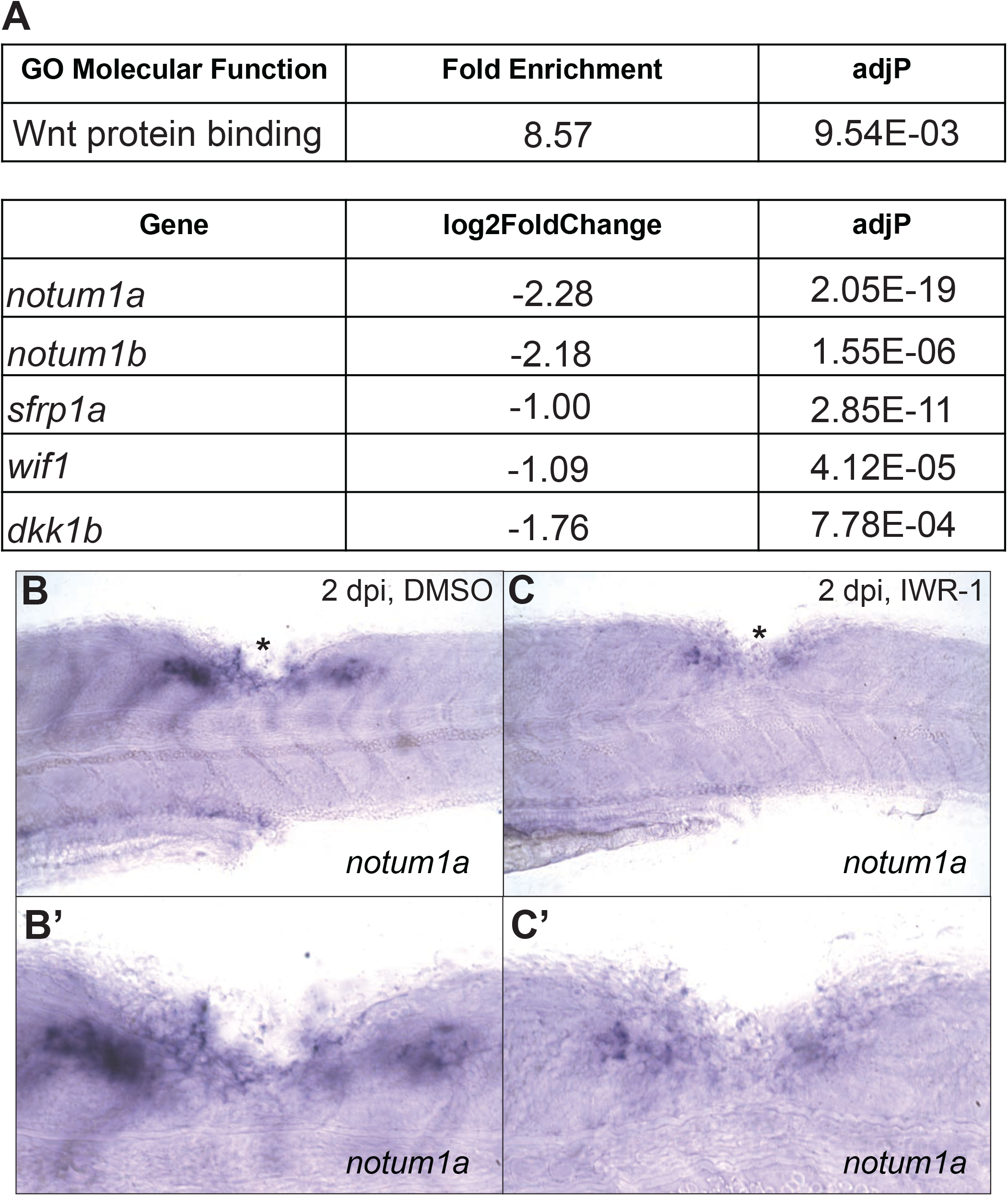
Wnt target genes are expressed in response to SCI. (A) Gene Ontology (GO) analysis of the 872 significantly downregulated genes from our RNAseq dataset shows significant enrichment of “Wnt protein binding”. IWR-1 treatment after SCI leads to significant downregulation of known Wnt target genes encoding secreted feedback inhibitors. (B, B’) *in situ* hybridization shows that *notum1a* is expressed in close proximity to the injury site at 2 dpi in control larvae. (C, C’) Injury-induced expression of *notum1a* is reduced by IWR-1 treatment. Asterisk: injury site.

### Wnt-dependent expression of infiltrating fibroblast markers

After SCI in larval zebrafish, most Wnt-responsive cells accumulate dorsal to the injury site ^10^. These cells have been classified as “fibroblast-like” based on tissue localization, collagen expression, and partial co-expression of the dermal fibroblast marker p63, but a more specific classification remains unknown ^10^. Furthermore, two *collagen12* paralogs are the only known Wnt-dependent genes expressed by fibroblasts and shown to be required for axon regrowth and functional recovery ^10^. Our RNAseq data showed significant downregulation of several transcripts known to be expressed by injury-responsive infiltrating fibroblasts, including orthologs of *fstl1, cthrc1*, and *ptx3* (Fig. 3A) ^21,27,28^. Our data also showed that Wnt inhibition leads to significant downregulation of two paralogous genes encoding the homeodomain transcription factor Prrx1, which has previously been implicated in fibroblast-mediated tissue regeneration (Fig. 3A) ^29-32^. In situ hybridization confirmed that IWR-1 treatment decreased the injury-induced expression of both paralogs, *prrx1a* and *prrx1b* (Fig. 3B-E’). A role for Prrx1 in central nervous system repair has not been described previously, and the Wnt-dependent expression of *prrx1a/b* after SCI could indicate a function for the pathway in the switch of infiltrating fibroblasts from a fibrotic to a proregenerative state, by regulating the expression of signals that promote neurogenesis and axon regrowth.

**Figure 3.**
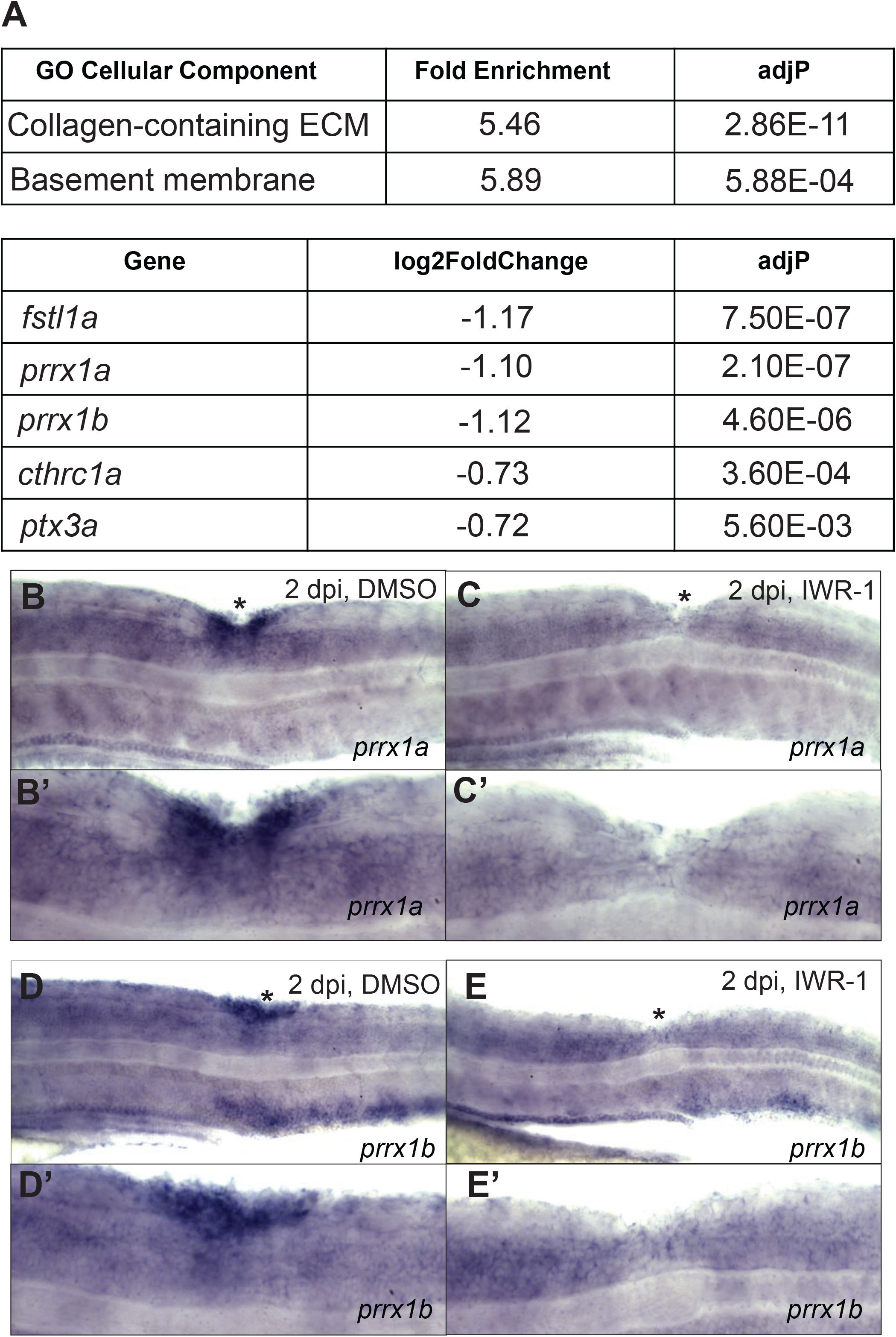
Infiltrating fibroblast markers are SCI- and Wnt-dependent. GO analysis of the 872 significantly downregulated genes from our RNAseq dataset shows significant enrichment in “basement membrane” and “collagen-containing ECM”. IWR-1 treatment after SCI leads to significant downregulation of transcripts previously reported to be expressed in injury-infiltrating fibroblasts. (B, B’; D, D’) *in situ* hybridization shows that *prrx1a* and *prrx1b* are expressed in close proximity to the injury site at 2 dpi in vehicle treated controls. (C, C’; E. E’) IWR-1 treatment reduces *prrx1a/b* expression after SCI. Asterisks: Injury site.

### Wnt-dependent expression of meningeal fibroblast markers

Meningeal fibroblasts are hypothesized to inhibit axon regrowth after mammalian SCI by secreting ECM components that contribute to chronic scarring and inflammation ^33^. In axolotls, which exhibit spontaneous spinal cord regeneration, putative meningeal fibroblasts aggregate on the leading edge of regrowing axons, consistent with a growth-promoting function ^34^. However, a meningeal role in zebrafish spinal cord regeneration has not yet been demonstrated. A recent study performed scRNAseq after injury to the mouse spinal cord and used two well-known meningeal cell markers, *decorin* and *col1a1*, to identify additional genes expressed in injury-responsive meningeal cells ^35^. We found that nine zebrafish orthologs of these 20 meningeal genes had significant Wnt-dependent expression in our RNAseq dataset. In addition, six of these genes, *fmoda, bambia, ogna, col1a1a, col1a1b*, and *aldh1a2*, show enriched expression in meningeal fibroblasts and meningeal precursor cells from Daniocell, an scRNAseq database of 0-5 dpf zebrafish, (Fig. 4A) ^36^, as well as orthologous enrichment in a scRNAseq dataset of embryonic mouse dural and arachnoid fibroblasts ^37^. These observations are also consistent with a previous report of Wnt activity in cells immediately adjacent to the spinal cord after zebrafish SCI ^10^, and thus suggest that Wnt-responsive meningeal fibroblasts may play a role in zebrafish spinal cord regeneration. While the cerebral meninges have begun to be characterized in zebrafish ^38^, the molecular and cellular composition of the spinal cord meninges and its roles in development, homeostasis, and regeneration remain poorly understood. We therefore sought to validate two of our RNAseq-identified candidate meningeal cell markers, *col1a1a* and *fmoda*. Expression was upregulated near the injury site at 2 dpi in controls, while treatment with IWR-1 after SCI decreased the expression of both transcripts (Fig. 4B-E’).

**Figure 4.**
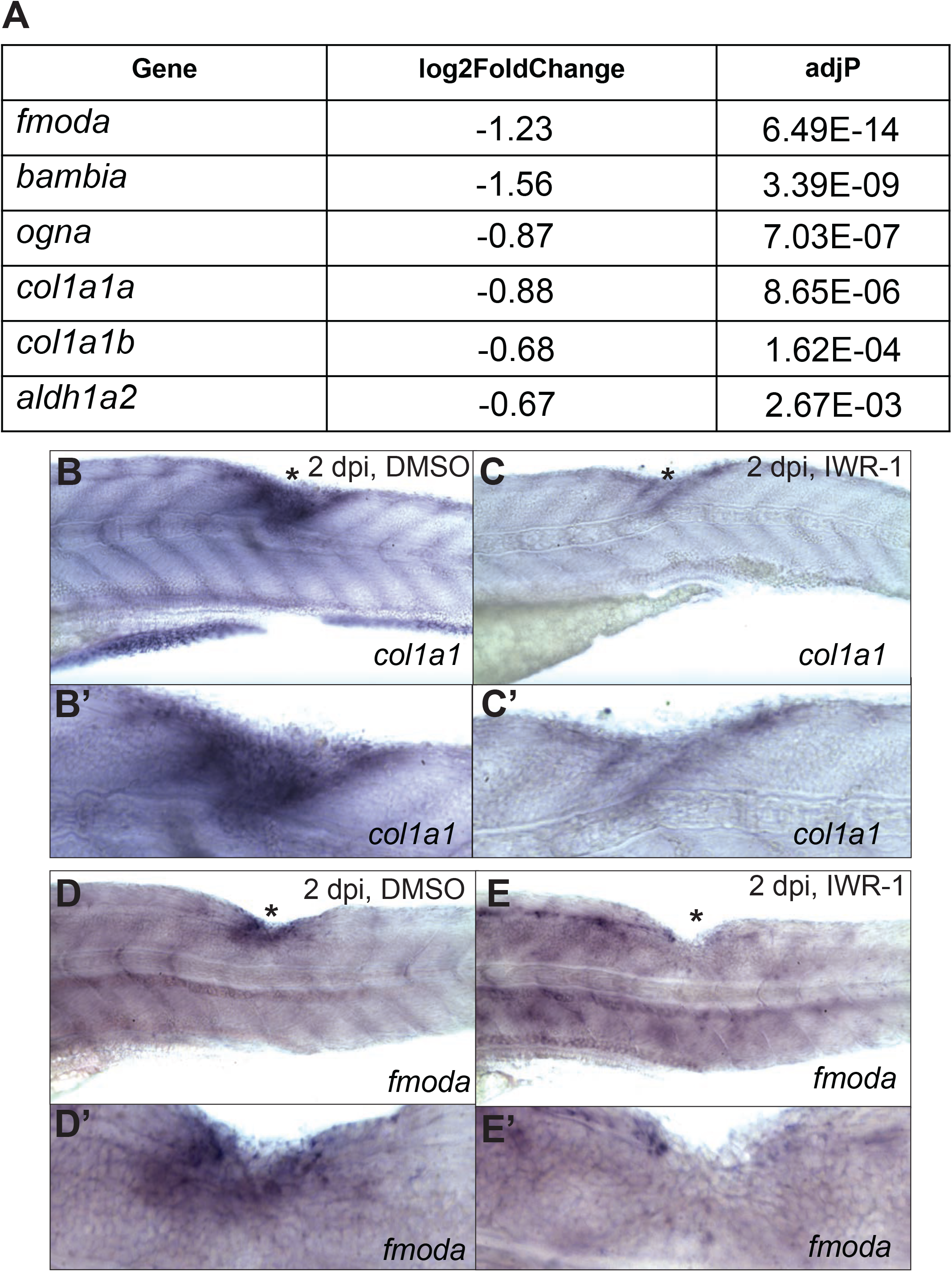
Markers for meningeal fibroblasts are SCI- and Wnt-dependent. IWR-1 treatment after SCI leads to significant reduction in the expression of markers known to be expressed in meningeal fibroblasts. (B, B’; D, D’) Two of these markers, *col1a1a* and *fmoda*, are expressed near the injury site in control larvae at 2dpi. (C, C’; E, E’) IWR-1 treatment reduces the SCI-induced expression of *col1a1a* and *fmoda* at 2dpi. Asterisks: Injury site.

Using a similar injury paradigm, Wnt signaling was previously shown to be required for *col12a* and *col12b* expression, but not for *col1a1a* expression, at 1 dpi ^10^. In that study, Wnt-independent *col12a* and *col12b* expression at 2 dpi was proposed to arise from a separate infiltrating fibroblast-like population through a relay signaling mechanism. Our observation of Wnt-dependent *col1a1a* expression at 2 dpi (Fig. 4B-C’) is also consistent with multiple waves of distinct Wnt-responsive cell populations appearing at the injury site over time. In addition, a recent study determined that *fmoda* expression is not increased, but in fact decreased at 1 dpi ^39^, while we observe clear upregulation by 2 dpi (Fig. 4D, D’). Together these results suggest that Wnt-responsive meningeal fibroblasts may repopulate the injury site after their initial loss following SCI. However, the extent to which this cell type contributes to spinal cord regeneration in zebrafish requires further investigation.

## CONCLUSIONS

In summary, our identification and validation of Wnt-dependent gene expression after SCI suggests that the subsequent inhibition of Wnt signaling may play a role in spinal cord regeneration, and identifies infiltrating and resident fibroblast populations that could participate in the repair process. Future studies will characterize the functional relevance of these genes and cell types in axon regrowth, neurogenesis, and the recovery of sensorimotor function.

## CONTRIBUTIONS

S.A.R. (Investigation, analysis, writing)

D.V. (Investigation, analysis, writing)

M.K.W. (Investigation, analysis, writing)

S.P. (Investigation)

R.I.D. (Funding acquisition, analysis, writing)

